# A Metabolic Reaction Balancing Web Service for Computational Systems Biology

**DOI:** 10.1101/187328

**Authors:** Paul D. Dobson, Pedro Mendes, Douglas B. Kell, Neil Swainston

**Affiliations:** School of Computer Science, The University of Manchester, Manchester, UK; School of Chemistry, The University of Manchester, Manchester, UK; Manchester Centre for Synthetic Biology of Fine and Speciality Chemicals (SYNBIOCHEM), Manchester Institute of Biotechnology, The University of Manchester, Manchester, UK; National Centre for Text Mining, The University of Manchester, Manchester, UK; Centre for Quantitative Medicine, School of Medicine, University of Connecticut, Connecticut, USA

**Keywords:** Systems Biology, metabolic modelling, reaction balancing, web service, CHO

## Abstract

**Background:** In metabolic network reconstruction the stoichiometric balancing of reactions is essential to create realistic constraint-based models. At the genome scale, balancing is a repetitive task that consumes valuable curator resource that could be deployed elsewhere. Automatic reaction balancing is possible and could be useful across computational systems biology, but widespread use of the appropriate code has been limited by the diversity of non-interoperable programming languages used in the field. RESTful web services offer a language-agnostic way of binding services together.

**Results:** Reaction balancing can be posed as a mixed integer linear programming problem to identify stoichiometric coefficients and infer commonly missing components. This functionality has been exposed as a web service that consumes a list of reactions as JSON or SBML. The reaction balancing web service has been deployed at http://www.nactem.ac.uk/balancer. Code is available via Github. By way of demonstration the service has been applied to a Chinese hamster ovary cell metabolic reconstruction to bring a further 219 reactions into balance.

**Conclusions:** The majority of systems biology software cannot access existing automatic reaction balancing tools due to a lack of language-specific bindings. Web services bridge different languages by using widely-spoken web communication protocols, meaning that one binding works for almost all languages. Automatic reaction balancing can now be consumed by any systems biology software via a RESTful web service.

## Background

In computational systems biology, metabolic network reconstruction is the long and arduous task of cataloguing all known small molecule reactions encoded by a genome [1]. One critical reconstruction subtask is to balance reactions by calculating stoichiometric coefficients. Within constraint-based models imbalanced reactions have the potential to create or consume mass unchecked by chemical constraints. This can lead to unrealistic behaviours as models exploit mathematical faults to create nutrients or dump excesses without regard for boundary conditions or biological reality. Realistic models cannot include unbalanced reactions.

Given reactant and product formulae and charges, it is generally not difficult to balance most reactions, but when approached manually at genome-scale this can consume considerable curator resource that might otherwise be deployed more gainfully. Some systems biology toolboxes [2, 3] implement tools to check whether or not a reaction is balanced, and even to bring protons into balance, but to the best of our knowledge automatic calculation of stoichiometric coefficients is only available within the Java-based SuBliMinaL Toolbox [4]. SuBliMinaL poses automatic reaction balancing as a mixed integer linear programming (MILP) that defines constraints across the reaction for each element and over charges. A MILP solver then adjusts stoichiometric coefficients in an attempt to satisfy these constraints. The balancing problem is complicated still further if chemical components have been inadvertently omitted. To address this common occurrence, SuBliMinaL also infers missing water and protons.

This ability to calculate stoichiometries automatically and even to infer missing components should have very broad application across computational systems biology, but being Java-based restricts the set of potential users. Rather than write custom bindings or re-implementations, a technology-agnostic binding like REST should be used. Here a Python re-implementation [5], written to bring balancing to Python (and therefore illustrative of the types of barrier we discuss), has been wrapped as a web service to bring automatic balancing to the widest possible audience. Below, the underpinning balancing mathematics are rehearsed, the web service interface is described with bindings to popular languages and toolboxes, and results from applying the service to a CHO metabolic reconstruction are shown.

### Implementation

In the SuBliMinaL Toolbox a reaction is represented within a mixed integer linear program by its right-and left-hand side components’ chemical formulae and charges, which must be brought into balance by modifying integer stoichiometric coefficients. Each chemical element in a reaction is represented as a balance constraint. To illustrate, consider the gross photosynthetic reaction scheme below. Three elemental balances are expressed for constituent elements C, H and O. The solver seeks positive integers for stoichiometric coefficients *R*_*1*_, *R*_*2*_, *P*_*1*_ and *P*_*2*_.

Reaction:

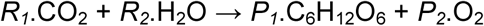

Constraint set:

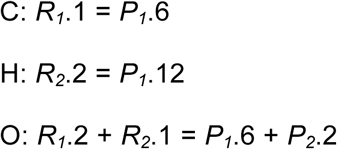

Charge is similarly incorporated but with the addition of signs. This defines a solution space for a Mixed Integer Linear Program (MILP). Optimal stoichiometric coefficients are found by minimising their sum.

Failure to balance a reaction properly is often attributable to the omission of commonly overlooked water or protons. These are incorporated automatically by their addition to each side of the reaction. The solver automatically eliminates these extra components again if they are not required. To illustrate, consider the incomplete version of the gross photosynthetic reaction:

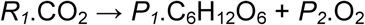

After adding water to both sides as a frequently missing component the reaction becomes:

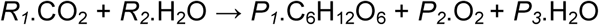

And the new constraint set becomes:

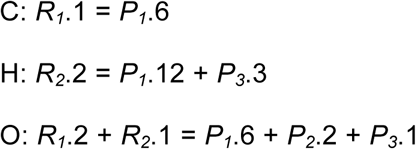

The solver identifies that *R*_*1*_, *R*_*2*_ and *P*_*2*_ = 6, *P*_*1*_ = 1 and, crucially, *P*_*3*_ = 0, correctly eliminating the unnecessary water from the right-hand side.

The automatic reaction balancing method above is available as a component of the Java-based SuBliMinaL Toolbox. The balancing function has since been re-implemented in Python to address some of the accessibility restrictions described earlier. To expose the balancer as widely as possible, the Python code has been wrapped as a web service using Flask-RESTful [6]. The underlying linear solver is GLPK [7] accessed via PyGLPK. For ease of installation, particularly as PyGLPK setup can be rather idiosyncratic on different operating systems, the web service runs within a Docker container based upon the Python 2.7 image. If very many reactions need balancing, this container should be used to set up a local web service in order to reduce hosting load. For smaller tasks the live web service can be accessed at http://www.nactem.ac.uk/balancer. Full code is available at https://github.com/dbkgroup/reaction-balancer.

The web service set up at http://www.nactem.ac.uk/balancer has three endpoints. A human endpoint is available at the landing page. Here a brief overview of the services, links to Github instructions and a trial interface are provided for manual use. Two web service endpoints: the JSON endpoint at http://www.nactem.ac.uk/balancer/balance/json and the SBML endpoint at http://www.nactem.ac.uk/balancer/balance/SBML. Technically only the JSON endpoint feeds into the elemental balancing MILP and solver as the SBML endpoint acts is a shim for converting SBML reactions to and from JSON for the JSON endpoint, although the user is hidden from this implementation detail.

The JSON endpoint expects a dictionary of uniquely identified reactions, sent via the POST request method. Each dictionary value specifies a reaction as a list of reaction participants. Each participant is specified using a list of four elements: formula, charge, current stoichiometry, with negatives denoting a reactant and positives denoting a product, and a reactant identifier, as in Figure 1.

**Figure 1.**
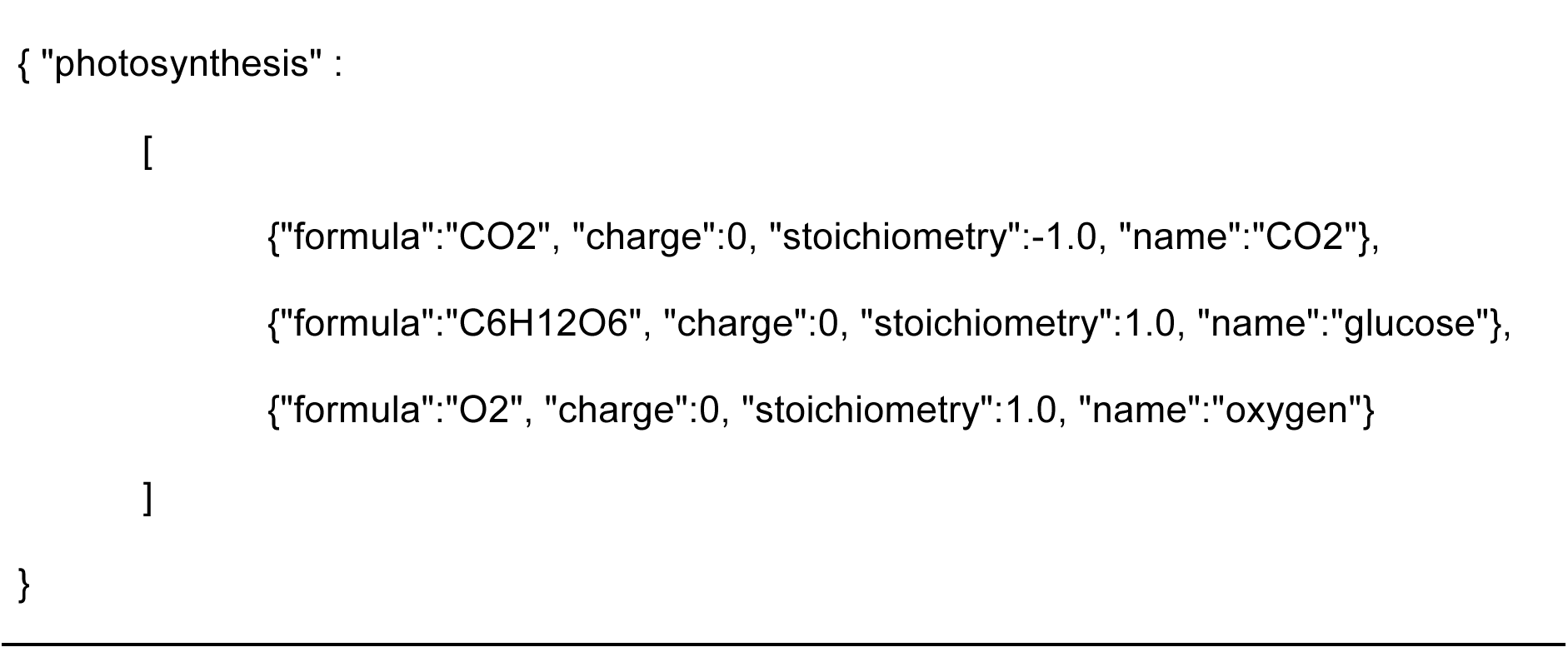
The JSON endpoint consumes a dictionary of reactions, with keys as unique reaction identifiers and values as lists of reaction components. Each reaction component is specified by chemical formula, charge, stoichiometry and an identifier.

The JSON response is similar to the input (Figure 2) but with dictionary values returning the reaction as the first component of a list, with booleans reporting whether or not the incoming (second element) and new reactions (third element) were balanced, and the final element reporting the status of the output. Where a reaction could not be balanced the returned reaction maintains its original stoichiometries, but otherwise is updated with the new stoichiometries and, where necessary, inferred components.

**Figure 2.**
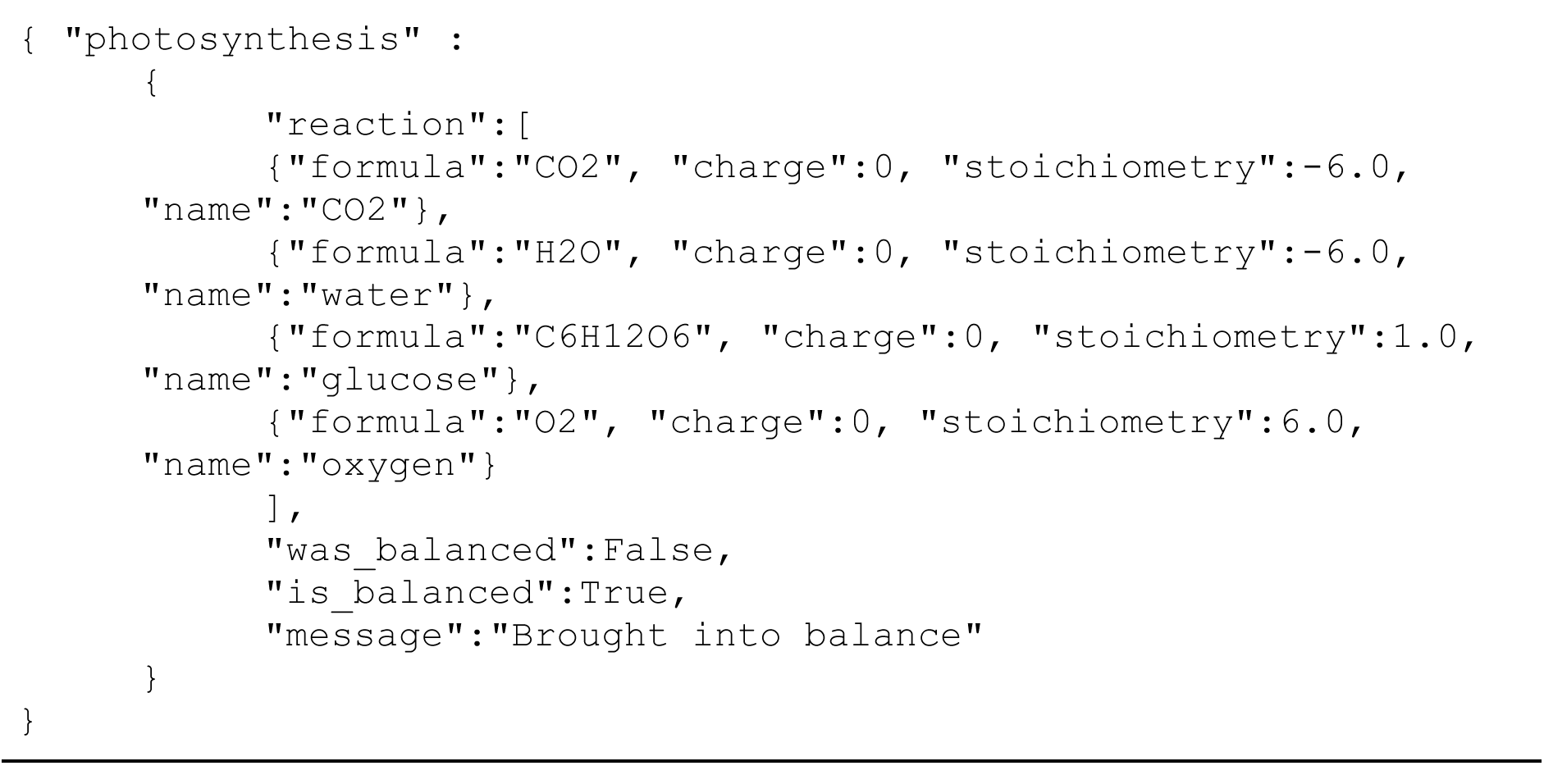
The JSON endpoint response is a dictionary of reactions, with keys as unique reaction identifiers and values that specify a list containing the updated reaction in the input format, a boolean reporting whether or not the input reaction was balanced, a boolean reporting whether or not the updated reaction is balanced, and a message summarising the response status. For reactions that could be assessed the message reports “already balanced” for reactions that were balanced on input, “brought into balance” for imbalanced reactions successfully brought into balance, and “not brought into balance” for imbalanced reactions not brought into balance. For reactions that could not be assessed because one or more components lacked formula and/or charge the message is “could not be assessed”.

The SBML endpoint expects SBML Level 3 [8] with the Flux Balance Constraints package [9]. To use the service with earlier versions of SBML users will need to write a custom parser to extract formulae and charges as, when present, these do not adhere to a standard representation. The SBML endpoint uses libSBML [10] to retrieve lists of species and reactions from the model. Reactions are checked to ensure all reactants and products have formulae and charges as the absence of either in any component means a reaction cannot be balanced. If all components are fully represented the reaction is converted to Figure 1 format and posted to the JSON endpoint. Responses are integrated into SBML with updated stoichiometries where applicable or original stoichiometries otherwise. The service output message is appended to the reaction Notes field.

When optional metabolites are added they are denoted with the suffix ‘_BALANCER’ as there is no simple way for optional metabolites to be mapped onto existing molecular species (species being distinct chemicals in localisations). Balance metabolites are added to the SBML list of species and located in a new compartment denoted ‘BALANCER’.

Use of the web service from different languages and toolboxes is demonstrated in the Examples folder of the Github repository. These include a generic Python binding using the requests package, cURL and Postman (a GUI for web API development).

## Discussion

Automatic reaction balancing with inference of commonly overlooked metabolites is, to the best of our knowledge, unique to the SuBliMinaL Toolbox. Related functions in other toolboxes check balances or adjust protons but do not go as far as calculating elemental balancing across the whole reaction to generate optimal stoichiometries. A web service interface to a Python implementation of SuBliMinaL’s automatic reaction balancer exposes this function in the most language-and platform-agnostic way possible. This makes automatic reaction balancing available to all computational systems biology software and, indeed, to the whole of chemistry.

The automatic reaction balancing service has been applied to the Chinese Hamster Ovary (CHO) metabolic network reconstruction iCHOv1 [1]. CHO cells are used for biopharmaceutical production as they are non-human (making virus transmission less likely) and able to synthesise, fold and glycosylate complicated protein products reproducibly at high titre. CHO-based therapeutic proteins are expensive because cell line development, which relies heavily upon labour-intensive and expensive screening and selection, and bioprocess regulatory approval, which demands exquisite control and reproducibility, cannot yet be designed in a Right First Time manner [11]. Genome-scale metabolic network reconstructions are being developed to enable computer-based design of optimal media and feeding strategies, metabolic engineering, and so forth. For metabolic models to drive cell factory design it is imperative that the reconstructions upon which they are built only describe balanced reactions. The iCHOv1 metabolic network reconstruction contains 4456 molecular species and 6663 reactions. Molecular species are specified using the fbc extension so the balancer was able to assess all reactions (although see comment below). 5316 reactions were already balanced in iCHOv1. A further 219 reactions were brought into balance by the automatic reaction balancing service using the SBML endpoint. 1128 reactions were not brought into balance. Of these, 637 were exchange reactions with only one component and therefore not intended to balance (they have been added for modelling purposes). The outstanding 491 reactions could not be balanced because, although correctly marked up, the formula and charge fields contained chemically inviable values (mostly unknown formulae denoted as “X” or blank, or formulae containing “R” groups). While there remains work to be done to bring iCHOv1 into full balance, for a version 1 release the reconstruction contained very few imbalanced reactions, due in no small part to efforts to capture formulae and charges fully. The automatic balancer was able to bring 219/5535 (4%) more of the fully specified reactions into balance. An updated iCHOv1 SBML is available in the Supplementary files (S2. iCHOv1.1.xml).

## Conclusions

Stoichiometric reaction balancing is a common task in metabolic network reconstruction, wider metabolic modelling and indeed across chemistry. Usually stoichiometries are simple enough to calculate manually but when undertaken at genome scale balancing becomes time-consuming, boring, and wasteful of expert resource. Automatic balancing is possible within the SuBliMinaL Toolbox but difficulties consuming this function within programming languages other than Java, and even with installing toolbox dependencies, have so far limited widespread uptake. The reaction balancing components of the SuBliMinaL Toolbox have therefore been written into a language-agnostic web service that is readily consumed by most modern programming languages.

The logical extension of exposing core functions as services is the microservice architectural style, which extends this strategy to build entire applications from loosely coupled web services. Microservice advocates highlight many useful properties of the style [12]. Of most relevance here is the ability of microserviced applications to mix languages, databases, operating systems and even hardware, as this provides a mechanism to bridge the centrally-uncoordinated, and therefore rather balkanised software that is typical in computational systems biology.

### List of abbreviations

MILP: Mixed Integer Linear Program

SBML: Systems Biology Markup Language

GLPK: GNU Linear Programming Kit

## Declarations

### Ethics approval and consent to participate

Not applicable

### Consent for publication

Not applicable

### Availability of data and material

All code, installation and operating instructions are available from Github at https://github.com/dbkgroup/reaction-balancer.

### Authors’ contributions

PDD wrote and deployed the web service, and wrote the manuscript. NS converted the original SuBliMinaL reaction balancing code to Python (subliminal-py) and wrote the core Dockerfile to run GLPK. All authors reviewed and contributed to the manuscript.

## Acknowledgements

The authors would like to acknowledge Paul and Ada Watson for useful advice on software design and deployment.

## Supplementary Files

S1. Gzipped archive of the Github code at time of submission.

S2. The Chinese hamster ovary cell metabolic network reconstruction with addition automatic balancing. iCHOv1_balanced.xml

